# Gene edited fluorescent cerebral organoids to study human brain function and disease

**DOI:** 10.1101/2020.11.24.395533

**Authors:** Lisa Bachmann, Lucia Gallego Villarejo, Natalie Heinen, David Marks, Thorsten Müller

## Abstract

Cerebral organoids are a promising model to study human brain function and disease, though the high inter-organoid variability of the mini-brains is still challenging. To overcome this limitation, we introduce the method of labeled mixed organoids generated from two different hiPSC lines, which enables the identification of cells from different origin within a single organoid. The method combines a gene editing workflow and subsequent organoid differentiation and offers a unique tool to study gene function in a complex human 3D tissue-like model. Using a CRISPR/Cas9 gene editing approach, different fluorescent proteins were fused to β-actin or lamin B1 in hiPSCs and subsequently used as a marker to identify each cell line. Mixtures of differently edited cells were seeded to induce embryoid body formation and cerebral organoid differentiation. As a consequence, the development of the 3D tissue was detectable by live confocal fluorescence microscopy and immunofluorescence staining in fixed samples. Analysis of mixed organoids allowed the identification and examination of specifically labeled cells in the organoid that belong to each of the two hiPSC donor lines. We demonstrate that a direct comparison of the individual cells is possible by having the edited and the control (or the two differentially labeled) cells within the same organoid, and thus the mixed organoids overcome the inter-organoid inhomogeneity limitations. The approach aims to pave the way for the reliable analysis of human genetic disorders by the use of organoids and to fundamentally understand the molecular mechanisms underlying pathological conditions.

## Introduction

Cerebral organoids (CO) offer a state-of-the-art 3D cell culture model to study human brain function and pathophysiology. COs are differentiated from human induced pluripotent stem cells (hiPSC) and comprise features of the human brain in respect to their cytoarchitecture, e.g. the different layers of the human brain, from the ventricular layer to the cortical plate. The protocol including the formation of embryoid bodies from hiPSC and the ectoderm-derived differentiation, which has been originally established in 2014 **(Lancaster and Knoblich, 2014)**, has meanwhile been extended to a variety of instructions to develop organoids containing specific brain areas like tissue with hippocampal neurons **(Qian et al., 2019)**. The applications of cerebral organoids for the basic research are increasing, which is a consequence of the Yamanaka protocol for the reprogramming of human body cells like PBMCs or fibroblasts into hiPSCs **(Takahashi and Yamanaka, 2006)**. As a consequence, hiPSCs can be obtained from every individual and similarly, it is possible to differentiate personalized cerebral (and other) organoids. COs could be on their way to become an indispensable technology for drug screening experiments **(Durens et al., 2020)**, biomarker identification, and drug treatment response analyses **(Salick et al., 2020)** for different fields of research, including neurodevelopment, neurodegeneration, and psychiatric disorders and the list is continuously growing. Despite these amazing features, the cerebral organoid technology is currently facing challenges, e.g. the lack of a co-differentiating vascular system (from the mesoderm germ layer), the absence of body axis, and a large heterogeneity in respect to their shape and size. As a consequence, many replicates need to be studied in order to define significant differences between patient-derived COs and controls 27402594 (Kelava and Lancaster, 2016).

Another technology of the 21th century is the nobel prize awarded CRISPR/Cas gene editing method offering unique capabilities to modify the genome of an organism. Originally identified as an adaptive immunity defense mechanism **(Barrangou et al., 2007)**, the technology is nowadays used to knock-out genes, to include mutations, or to activate and silence genes **(Bukhari and Müller, 2019)**. Although it has been mainly used for the aforementioned purposes, CRISPR/Cas has already been pointed out as a promising tool for future applications in cell tracing and reporter-cell generation *via* specific expression of fluorescent proteins **(Fischer et al., 2019), (Bagley et al., 2017)**. The application of CRISPR/Cas gene editing on hiPSCs and the subsequent 3D organoid differentiation promise a fascinating tool to study the consequences of genetic changes in a 3D tissue-like model and provide an auspicious technology to examine the molecular mechanisms underlying human physiology and pathophysiology.

Accordingly, in the current study, we focused on an innovative way to combine two outstanding molecular biological techniques, CRISPR/Cas9 gene editing and cerebral organoid generation, in order to overcome some of the limitations accompanying CO technology. We used fluorescent-labeled cell lines, tagged for broadly expressed proteins with a fluorescent cassette. Using this approach, we generated functional cerebral organoids that allow the identification of differently labeled cells by confocal microscopy. We further observed a comparable organization and cell type composition in fluorescently labeled COs as reported for non-edited COs. Moreover, our results confirm the possibility to develop “mixed” cerebral organoids from gene-edited cells with different genetic backgrounds and different labels that can be identified and analyzed separately within the same CO. Altogether, our results support the possibility of using distinctively labeled reporter cell lines, with different characteristics (e.g. mutation-containing and “healthy cells”) to generate more accurate human brain disease models enabling a direct comparison under identical experimental conditions.

## Material and Methods

### HiPSC culture

HiPSCs were cultured in StemFlex™ medium (Gibco™) with 1 % Penicillin/Streptomycin on Geltrex™ (Gibco™, 1:150 dilution in DMEM/F12) coated 35 mm dishes at 37 °C in 5 % CO_2_ atmosphere. The parafilm-sealed dishes were stored at 4 °C for up to 2 weeks and were incubated for 60 min at 37 °C prior to use for polymerization. A light microscope was used to observe cell growth and health. The StemFlex™ medium was changed every 1-2 days and the cells were split before 70-80 % confluency was reached. For splitting, the cells were rinsed with 1 ml DPBS (Gibco™) once and harvested by adding 1 ml ReLeSR™ (Stem Cell) for 30 s at RT. The solution was removed except for a thin film and incubated for additional 90 s, occasionally tapping the dish gently on the bench. The undifferentiated cell colonies were detached by rinsing 3-4 times with 1 ml StemFlex™ and divided on new culture dishes.

### Gene editing of HiPSC

Before 70 % confluence was reached, TrypLE™ (Gibco™) was used to detach the cells according to the manufacturer’s instructions. The cells were counted using a Neubauer chamber and 200.000 cells per nucleofection sample were prepared according to the manufacturer’s instruction using a Nucleocuvette (Lonza) and P3 primary cell nucleofection buffer (Lonza). Nucleofection was performed according to the instructions of IDT with Nucleofection enhancer, RNP complex (Cas9 with tracrRNA for LMNB1 gene) (IDT) and 400 ng of LMNB1_mTagRFP-T HDR plasmid (Addgene 114403). Nucleofection was done with the pulses DN-100, CA-137 and control (no pulse) with the Lonza 4D nucleofector.

HiPSCs were cultivated with 0.1 % ROCK inhibitor (Stem Cell) and 0.1 % HDR enhancer at 37 °C and 5 % CO_2_ and the medium was changed 24 h after nucleofection. The fluorescent signal of the knock-in (KI) of the cells was observed by fluorescence microscopy 72 h after nucleofection. Several days post nucleofection, cells were detached using TrypLE™ (Gibco™) as described before and centrifuged. The pellet was resuspended in DPBS in a concentration below 100.000 cells/ml. Cells that successfully included the RFP fluorescent KI were separated by fluorescent-assisted cell sorting (FACS) using the MoFlo Astrios Eq Cell Sorter (Beckman Coulter). More than 3000 cells were sorted in one well of a 96-well plate pre-coated with Geltrex™ and equilibrated with 100 μl Stemflex™ and 0.1 μl ROCK inhibitor. Cells were further incubated at 37 °C and 5 % CO_2_ and cultured changing medium every other day. The ACTB-GFP hiPSC line is commercially available and was purchased from Coriell Institute (AICS-0016 hiPSC).

### Normal and Mixed Cerebral Organoids

Cerebral organoids were cultured according to the protocol of Lancaster and Knoblich **(Lancaster and Knoblich, 2014)** and only modifications are stated in the following. HiPSCs were harvested using TrypLE™ and hiPSC lines (ACTB-GFP, RFP-LMNB1, CL1, CL2 and CL3) were mixed in respective ratios before centrifugation at 300 x g for 5 min at RT. On day 1 and 2 after seeding, the EBs were monitored for their appearance and size. If the EBs showed blurred edges or a diameter smaller than 300 μm, half of the medium was aspirated and exchanged with 150 μl of fresh hESC medium containing 4 ng/ml bFGF and 50 μM ROCK inhibitor. On day 3, the medium was changed to hESC medium without supplements. After six days (day 6), the medium was changed to NI medium and then changed every day until day 11 or 12, when EBs were embedded in growth-factor reduced Matrigel (Corning) droplets. Therefore, the EBs were removed from the well using a cut 200 μl pipette tip and placed in a trough on a parafilm strip. Excess media around the EBs was removed and a 10 μl drop of Matrigel was added to each EB which was then carefully moved to the middle of the drop using a 10 μl pipette tip. After 25-30 min of incubation at 37 °C to allow polymerization of the Matrigel, the Matrigel drops were carefully loosened at the edges using a 10 μl pipette tip and the parafilm strip was placed upside down into a well of a six-well plate filled with 2 ml of DM-A medium. Gentle shaking of the plate detached the Matrigel drops from the parafilm strip and the parafilm strip could be removed from the well. Embedded EBs were kept at 37 °C and 5 % CO_2_ in a humidified atmosphere.

### Organoid cryo- and vibratome-sections and immunofluorescence staining

COs were fixated in 4 % PFA (4 °C, 90 min) and subsequently washed and incubated in a 30 % sucrose solution (4 °C, overnight). The next day, the COs were embedded in a 1:1 mixture of 30 % sucrose solution and Tissue freezing medium (Sakura), snap-frozen on dry ice, and stored at −80 °C. They were sliced into 16 μm sections using a cryostat (Leica CM3050S), mounted on Superfrost™ slides (Thermo Fisher), and stored at −80 °C until further use. For immunofluorescence staining, the following steps were performed at RT and protected from light and washing was done with PBS for 4 min, if not stated otherwise. The CO sections were thawed in PBS and circled with a Liquid Blocker Super PAP pen (Thermo Fisher). After washing twice, the sections were blocked and permeabilized with 0.1 % Triton-X-100 and 5 % goat serum in PBS for 1 h. Primary antibodies were diluted in 0.1 % Triton X-100 in PBS (β - tubulin III: ab78078, abcam, 1:500; MAP2: M4403, Sigma, 1:500; PAX6: 60094, StemCell, 1:500) and incubated on the CO sections in a humidified chamber at 4 °C overnight. The sections were washed twice and incubated with the (biotinylated) secondary antibody, diluted in 0.1 % Triton-X-100 in PBS, in a humidified chamber for 1 h. After washing twice, the sections were incubated with Avidin-TRITC diluted 1:1000 in 0.1 % Triton X-100 in PBS. In case of not using a biotinylated antibody, the secondary antibody was diluted in 0.1 % Triton X-100 in PBS and incubated on the sections in a humidified chamber for 45 min. The sections were washed two times and incubated with 0.001 mg/ml DAPI or 0.001 mg/ml Hoechst33342 for 15 min. Following two washing steps, the sections were mounted with a coverslip and left for drying overnight.

Vibratome slices were generated according to the protocol by Lancaster et al. (Giandomenico et al., 2019) with some modifications. Briefly, mature organoids (day 35-45) were washed (PBS without Ca^2+^ and Mg^2+^) and embedded in 3 % low-melting-point agarose (Thermo Fisher Scientific) at 40 °C in reusable silicone molds (1 cm³). The agarose blocks were cooled on ice for 10 min and sectioned into 300 μm thick slides, using the Leica VT1000S vibrating microtome in cold PBS. Sections were collected in 24-well plates, containing serum-supplemented slice culture medium (SSSCM: DMEM (Thermo Fisher Scientific), 10 % FBS, 0.5 % (w/v) glucose, 1x (v/v) GlutaMAX (Thermo Fisher Scientific), 1 % Antibiotic-Antimycotic (Thermo Fisher Scientific) and were incubated for 1 h at 37 °C. After incubation, the medium was replaced with serum-free slice culture medium (SFSCM: Neurobasal (Thermo Fisher Scientific), 1:50 (v/v) B-27 supplement (Thermo Fisher Scientific), 0,5 % (w/v) glucose, 1x (v/v) GlutaMAX (Thermo Fisher Scientific) and 1 % Antibiotic-Antimycotic (Thermo Fisher Scientific). Sections were cultivated at 37 °C, 5 % CO2 and the medium was exchanged every other day.

### Widefield and confocal microscopy

Developing fluorescently labeled EBs were regularly imaged using an Olympus IX51 inverted microscope equipped with a 10x objective. Immunohistochemically stained CO sections were imaged with a Leica TCS SP8 confocal microscope system using a 10x dry objective (0.3 NA) or 100x oil objective (1.4 NA) and a 405 nm and white light laser with excitation wavelengths of 405 nm (DAPI/Hoechst), 488 nm (EGFP) and 551 nm (TRITC) and emission wavelengths of 461 nm (DAPI/Hoechst), 509 nm (EGFP) and 576 nm (TRITC). The detection ranges were 410-463 nm (DAPI/Hoechst), 493-592 nm (EGFP) and 556-701 nm (TRITC). Images were recorded in a sequential scan at a scan speed of 600 Hz into 5648 x 5648 (10x) or 2640 x 2640 (100x) images. Hybrid detectors (HyD) were used for EGFP/TRITC and a photomultiplier tube detector (PMT) for DAPI/Hoechst. For tile scans, 465 x 465 μm areas (3 x 3 tiles) were recorded in x- and y-direction. The LAS X-software was used to analyze the data, and to improve resolution, a deconvolution process was performed via HyVolution 2 in the Huygens Essential Automatic approach, taking the refractive index of the mounting medium into account.

## Results

### Setup of the mixed organoid approach using CRISPR/HDR labeling of hiPSC

Mixed organoids were derived from human induced pluripotent stem cells (hiPSC), which have been obtained by re-programing of patient derived fibroblasts or peripheral blood mononuclear cells (PBMCs, Figure 1A **(Takahashi et al., 2007)**). Within our approach we used the commercially available edited β-actin (ACTB-GFP) hiPSC line (Figure 1B). In addition, we labeled the Lamin B1 protein (LMNB1) and included a red fluorescent protein (RFP). To generate the latter, a CRISPR/Cas9 homology directed repair (HDR) approach was used and RFP was fused N-terminal to the LMNB1 gene as shown before **(Roberts et al., 2017)**. Upon fluorescence activated cell sorting (FACS), pure RFP-LMNB1 hiPSCs were obtained and subsequently used for cerebral organoid differentiation (Figure 1C). As an initial approach, we aimed to test the pluripotency of the edited cells to differentiate into cerebral organoids **(Lancaster and Knoblich, 2014)**. Successful differentiation would result in a labeled cerebral organoid with a GFP positive β-actin signal in the cytosol or an RFP positive outlined nucleus, respectively. Furthermore, mixtures of the fluorescently labeled cells were expected to result in cerebral organoids, in which each cell is traceable to its original hiPSC line (Figure 1C).

**Figure 1.**
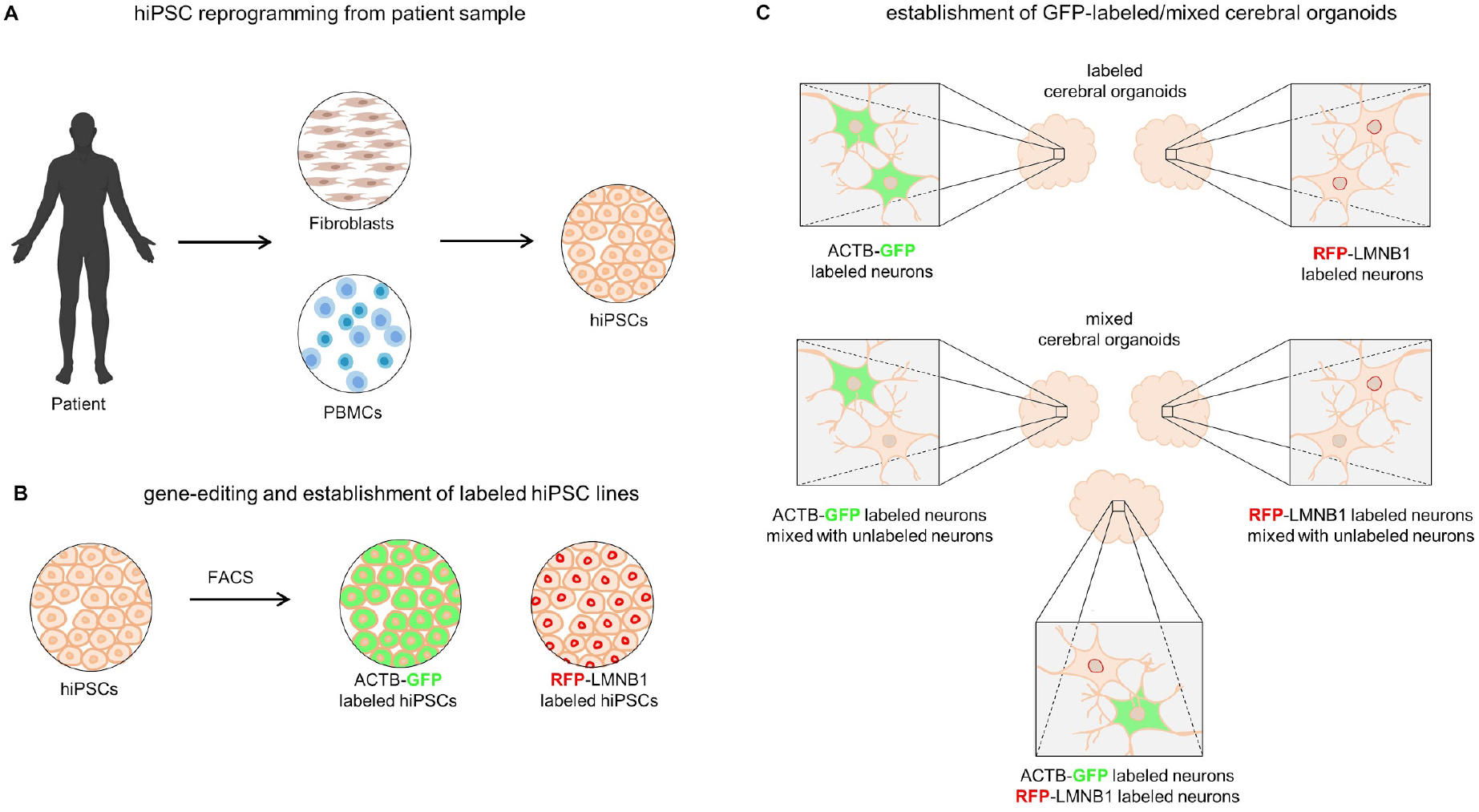
From patient-derived samples to gene-edited mixed cerebral organoids. **(A)** Somatic patient-derived cells like fibroblasts or peripheral blood mononuclear cells (PBMCs) can be reprogrammed to hiPSCs following the Yamanaka protocol. **(B)** HiPSCs can be used to knock-in a fluorophore with the CRISPR/Cas9 system. In this study, β-actin (ACTB) was C-terminally labeled with GFP; lamin B1 (LMNB1) was N-terminally labeled with RFP. The cells were purified with fluorescent activated cell sorting (FACS). **(C)** Labeled hiPSCs were used to generate cerebral organoids following the protocol of Lancaster et al.. Two different approaches were followed: organoids comprising only labeled cells (ACTB-GFP/RFP-LMNB1) or mixed organoids comprising labeled and unlabeled or differentially labeled hiPSCs.

### Gene-edited ACTB-GFP hiPSC differentiate into regular cerebral organoids

As an initial approach, we aimed to test the impact of the ~25 kD GFP fusion tag on the potential of hiPSC to establish cerebral organoids. After seeding, ACTB-GFP hiPSC demonstrated differentiation of embryoid bodies with sharp edges and a round shape (Figure 2A). On day 6, when neural induction is started, the EBs have grown visibly in size and have lost their completely round form. Upon the day of embedding (day 12) the formation of epithelial buds was observed, and the size and extent of budding increased in the subsequent weeks and an organized layering was evident (e.g. day 36). The endogenous ACTB-GFP labeling enabled us to study the development by fluorescence microscopy revealing an evenly distributed GFP signal (Figure 2B). Due to the increasing organoid diameter and a limited laser excitation penetration depth, fluorescence imaging was restricted to the first ~14 days, however preparation of living vibratome sections according to published protocols **(Giandomenico et al., 2019)** enabled the study of older organoids (sections) in live over time (Figure 2D). Live imaging revealed reorganization of some parts within the CO (video 1, Supplement 1). Within the whole period of differentiation, the size of the edited EBs/organoids was comparable to that of non-edited cell lines (CL1) with different genetic background (Figure 2C). Late ACTB-GFP organoids (> 30 days) tended to be of minor size than CL1 organoids but this difference failed to achieve significance. Fluorescence microscopy using higher resolution (100x) demonstrated the development of cerebral organoid sub-structures (Figure 2E). First two images show the development of a layered structure containing lumen, ventricular/subventricular zone and cortical plate (corresponding video 2 (z-scan) and video 3 (time-scan), Supplement). Third image demonstrates growth of the organoid layer by layer (video 4, Supplement). Last image demonstrates the β-actin related cyto-architecture of the living organoid in high resolution.

**Figure 2.**
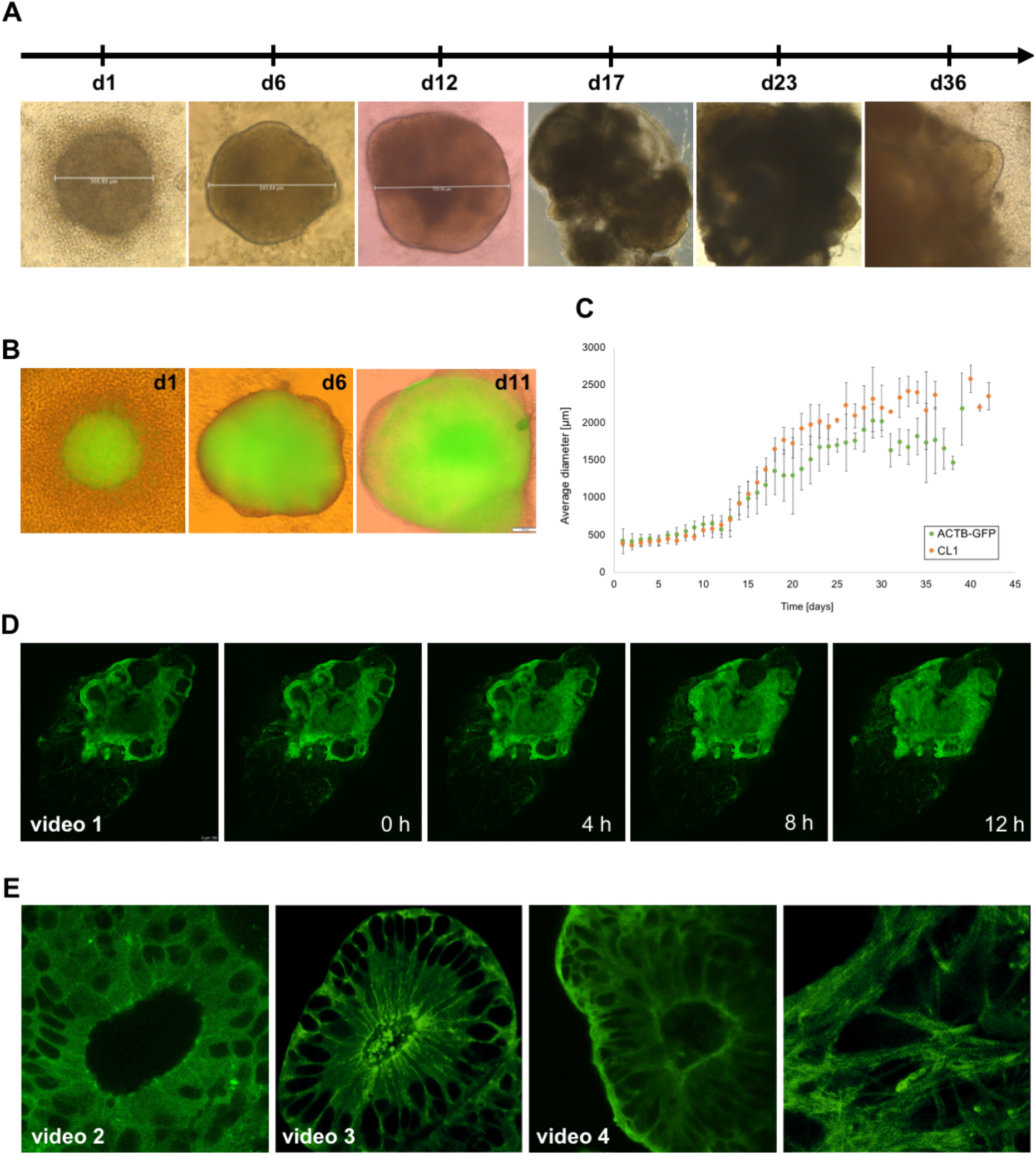
Differentiation of ACTB-GFP cerebral organoids. Gene-edited hiPSCs with an ACTB-GFP fluorescence label were seeded to form EBs and cultured to develop into cerebral organoids. **(A)** Light microscopy imaging revealed successful formation of EBs (d1) with cleared edges (d6), development of translucent ectoderm after neural induction (d12), and neuroepithelial budding after Matrigel embedding (d17, d23, d36). **(B)** Fluorescence microscopy imaging of whole EBs showed an evenly distributed, progressively intensifying fluorescence signal on d1, d6 and d11. **(C)** The average diameter over the course of cultivation time is depicted as a growth curve for ACTB-GFP EBs/COs in comparison with a non-edited CL1 EBs/COs. Errors bars represent standard errors from at least three biological replicates. EBs starting at a diameter of 400-500 μm grew steadily and exhibited extensive growth after Matrigel embedding. Growth tendentially (without significance) differed after around 25 days and at 2000 μm of size, when CL1 COs continued to increase and ACTB-GFP COs decreased in diameter. **(D)** Live confocal fluorescence imaging of vibratome-sliced COs revealed ongoing reorganization of the tissue (video 1 given in the supplement). **(E)** First two images (z-scan in video 2, time scan in video 3, supplement) demonstrate the live development of CO compartments including a central lumen, VZ/SVZ, and cortical plate. Third image (video 4) illustrates the CO growth layer by layer, and the last image (video 5) shows the structured ACTB-GFP signal in high resolution.

### Mixed Organoids of two different non-isogenic hiPSC grow and differentiate like monocellular organoids

Seeding of ACTB-GFP hiPSCs mixed with unlabeled hiPSC lines (CL1, CL2, CL3) in different ratios led to the formation of embryoid bodies with defined edges and a round shape comparable to monocellular derived organoids (Figure 3A, Supplementary Figure 1A). Compared with ACTB-GFP EBs, mixed EBs containing CL1 and CL2 had a less regular shape after seeding and exhibited a deformation of shape after neural induction (day 6). However, this did not affect successful development of the neuroectoderm, as apparent from translucent edges, or formation of neuroepithelial buds after Matrigel embedding (day 12). Similar to pure ACTB-GFP COs, an increase in size and extent of budding could be observed. The endogenous label was observable by fluorescence microscopy imaging of whole EBs seeded from 75 % ACTB-GFP and 25 % CL1/CL2 hiPSCs as well as by imaging of EBs seeded from 50 % ACTB-GFP and 50 % CL3 hiPSCs, whereby a difference to pure ACTB-GFP EBs was in particular evident in 50 % ACTB-GFP EBs revealing a more speckle-like distribution of the GFP marker which is only visible in 75 % ACTB-GFP EBs on d1 (Figure 3B, Supplementary Figure 1B). Mixed EBs showed progressive growth starting at ~500 μm in diameter with a rise after Matrigel embedding, and reached an average diameter of ~2500 μm after 40 d of cultivation (Figure 3C, Supplementary Figure 1C, shown for three different cell lines with different mixtures), which was in line with pure ACTB-GFP COs (Figure 2C) and unlabeled controls. A significant difference in size development between COs seeded from different ratios of ACTB-GFP and CL1/CL2/CL3 hiPSCs was not discernable (Supplementary Figure 1D).

**Figure 3.**
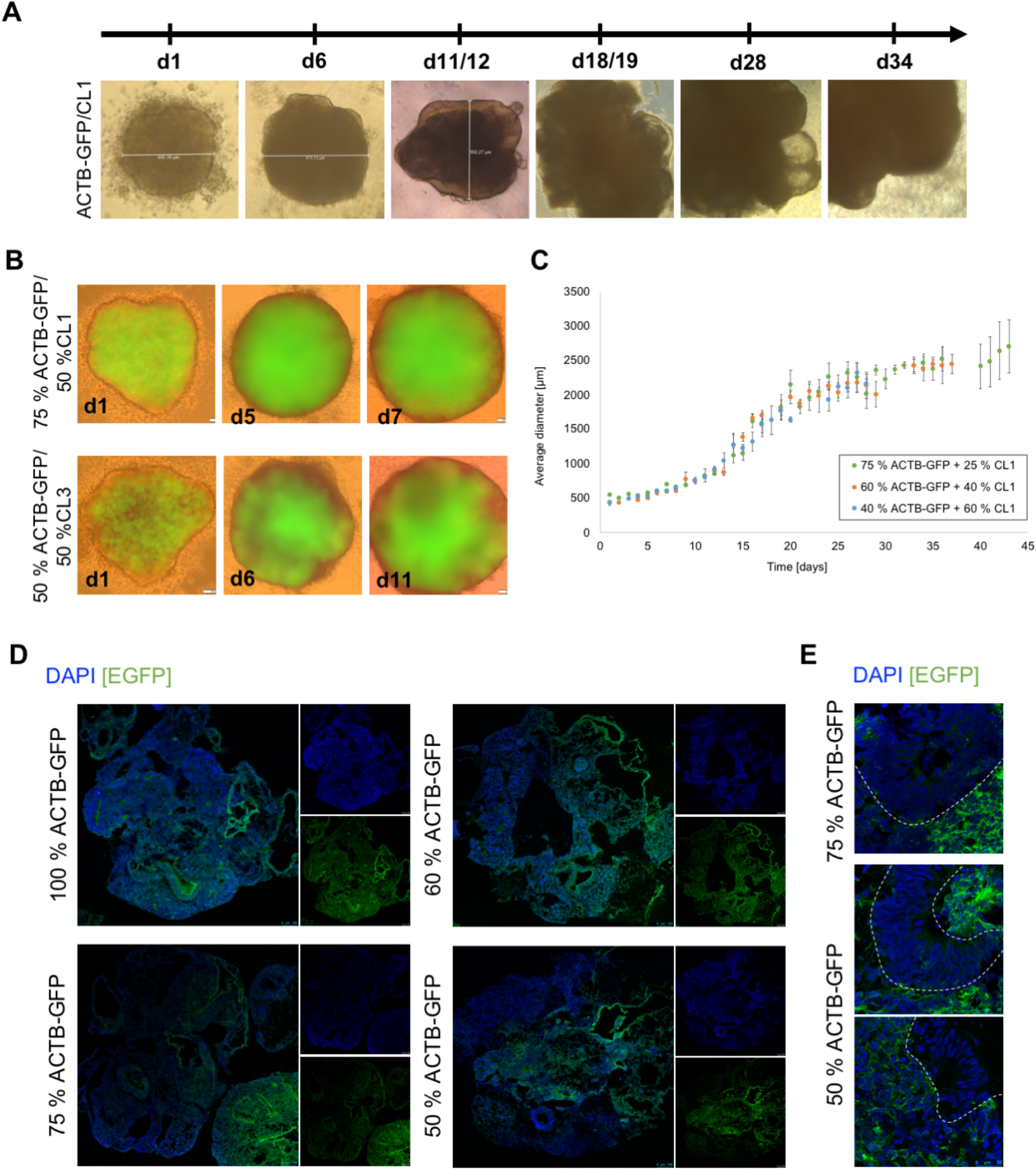
Development of mixed cerebral organoids. Cerebral organoids were generated from ACTB-GFP gene-edited hiPSCs mixed with different hiPSC lines (CL1, (CL2, CL3 in Supplementary Figure 1A)) in different ratios. **(A)** Light microscopy imaging revealed successful formation of EBs with defined edges and a round shape. Translucent edges and the formation of neuroepithelial buds after Matrigel embedding (day 12) was detectable for all mixtures. **(B)** Fluorescence microscopy demonstrated the endogenous label in EBs seeded from 75 % ACTB-GFP + 25 % CL1 hiPSCs. A speckle-like distribution was particularly evident in EBs seeded from 50 % ACTB-GFP + 50 % CL3 (ACTB-GFP/CL2 in Suppl. Fig. 1B). **(C)** Growth of mixed EBs (different mixtures of ACTB-GFP and CL1 (CL2/CL3 in Suppl. Fig. 1D)) starting at ~500 μm and reaching up to 2500 μm after 40 days of cultivation was not different to the growth of pure ACTB-GFP or unlabeled control samples. Error bars correspond to the standard error from at least three biological replicates. **(D)** Confocal microscopy imaging of COs stained for DAPI shows a reduction in GFP signal with decreasing percentage of ACTB-GFP in 10x magnification. **(E)** 100x magnification of 75 % and 50 % ACTB-GFP COs reveals structures with an uneven distribution of GFP signal.

After 28 d of differentiation, COs were fixed and sliced in order to assess the GFP distribution throughout the whole organoid in more detail (Figure 3D). 10x overview confocal fluorescence microscopy imaging revealed positive fluorescence of the whole 100 % ACTB-GFP organoid slice as expected. Imaging of organoids differentiated with decreasing ratios of ACTB-GFP cells demonstrated a reduction in the overall fluorescence intensity. High resolution imaging (100x objective) demonstrated a patterned distribution of the GFP signal in 75 % and 50 % mixed COs (Figure 3E) including GFP positive areas as well as unlabeled areas within the same organoid. As a consequence, each single cell is assignable to their origin, to the GFP labeled or the non-labeled hiPSC line.

### Mixed organoids demonstrate differentiation like monocellular organoids

As a next step, gene-edited monocellular and mixed cryo-sectioned COs were analyzed in respect to their cerebral differentiation. Immunohistochemical staining revealed tube-like structures including a β-tubulin III positive layer of immature neurons (Figure 4A) constituting a cortical-plate-like layer (CP), which encloses a PAX6 positive layer of radial glia (Figure 4B) forming the (sub-)ventricular zone (VZ/SVZ) and surrounds a central lumen (Lu). Similar cortical structures could be observed in pure ACTB-GFP COs as well as in mixed COs (Figure 4C,D). Additional staining against neuronal dendrite marker MAP2 showed the presence of mature neurons in the cortical plate-like layer surrounding the (sub-)ventricular zone and lumen in ACTB-GFP (Figure 4E) and in mixed COs (Figure 4F).

**Figure 4.**
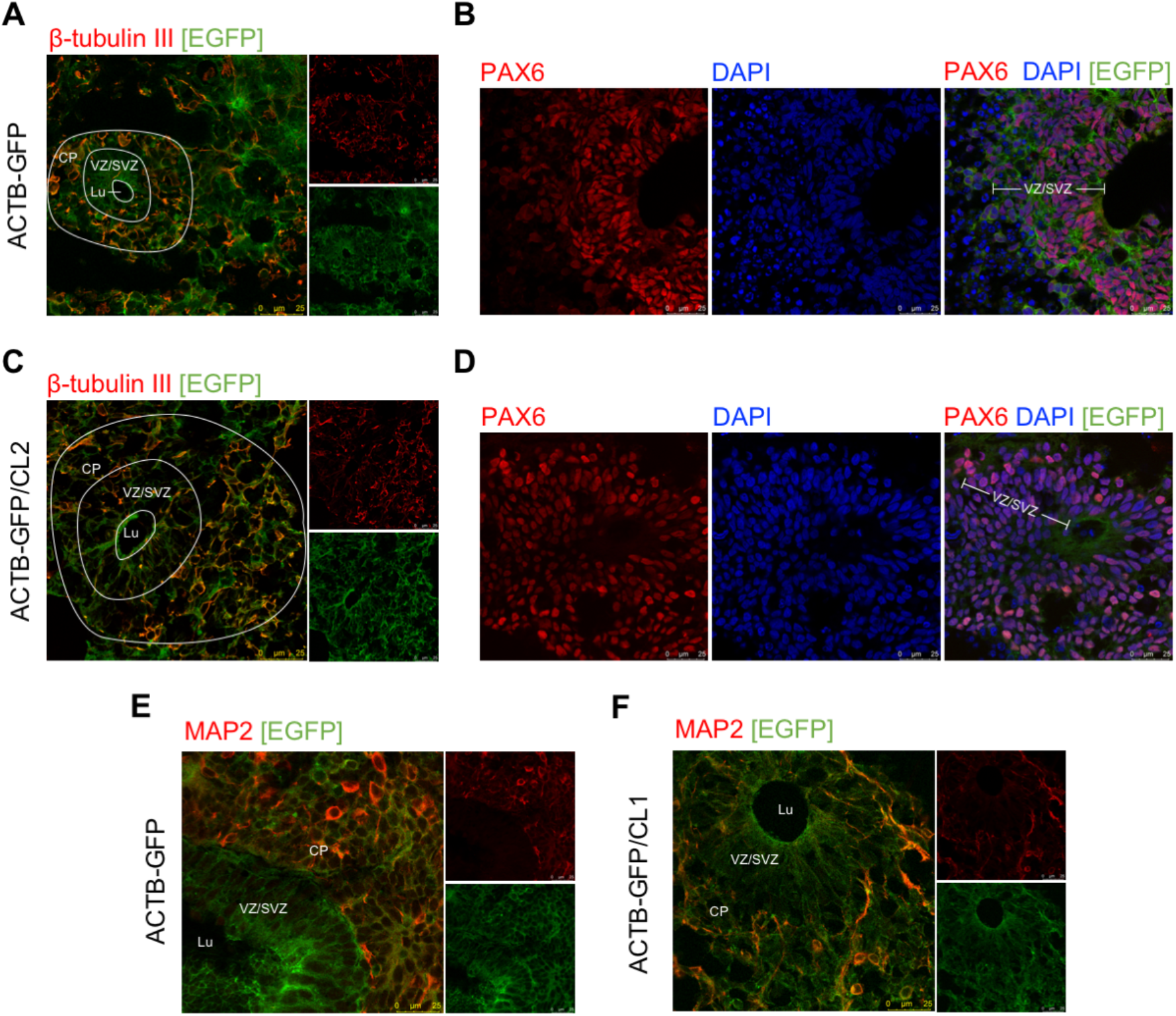
Differentiation of cortical structures in ACTB-GFP and mixed cerebral organoids. Cerebral organoids generated from ACTB-GFP hiPSCs or mixed hiPSC lines were cryosectioned and immunohistochemically stained for neuronal markers β-tubulin III **(A, C)** and MAP2 **(E,F)** as well as neural progenitor/radial glia marker PAX6 **(B, D)**. Tube-like structures containing a (sub-)ventricular zone (VZ/SVZ) of PAX6 positive radial glia surrounding the lumen (Lu) are encircled by a layer of β-tubulin III positive immature neurons or MAP2 positive mature neurons forming a cortical plate-like layer (CP).

### Proof of principle - Mixed organoids from RFP-LMNB1

In order to validate the mixed organoid approach we used the nuclear envelope protein Lamin B1 as further target and introduced an N-terminal RFP tag using a CRISPR/Cas9 HDR approach **(Roberts et al., 2017)**. Edited hiPSC showed a prominent staining of the nuclear envelope, were highly viable and proliferated like the parental line. The cell lines retained their pluripotency as shown by their potential to develop EBs being similar to other EBs in respect to their roundish shape with defined edges and translucent edges from the age of 8 days (Figure 5A). When analyzed by fluorescent microscopy, EBs differentiated from RFP-LMNB1 as well as EBs from 50 % RFP-LMNB1 and 50 % ACTB-GFP showed normal development with an apparent pattern-like distribution of the fluorescence labeled cells (Figure 5B). Moreover, confocal fluorescence analysis of EBs from 50 % RFP-LMNB1 and 50 % ACTB-GFP demonstrated the uneven distribution of the different cell lines (Figure 5C). Result have been confirmed with additional staining and confocal analysis (Figure 5D) demonstrating the generation of cell-type derived areas within one mixed EB (Figure 5E) comparable to the results obtained from the mixture including ACTB-GFP and non-labeled COs (Figure 3E). High resolution imaging of CO slices (Figure 5E) revealed two populations: cells derived from the RFP-LMNB1 hiPSC with a distinct red labeling of the nuclear envelope and cell pattern derived from the ACTB-GFP hiPSCs with green cytoplasmic fluorescence. These results together highlight the feasibility of the mixed organoid approach enabling the comparison of two different hiPSC derived cell pattern within the very same organoid.

**Figure 5.**
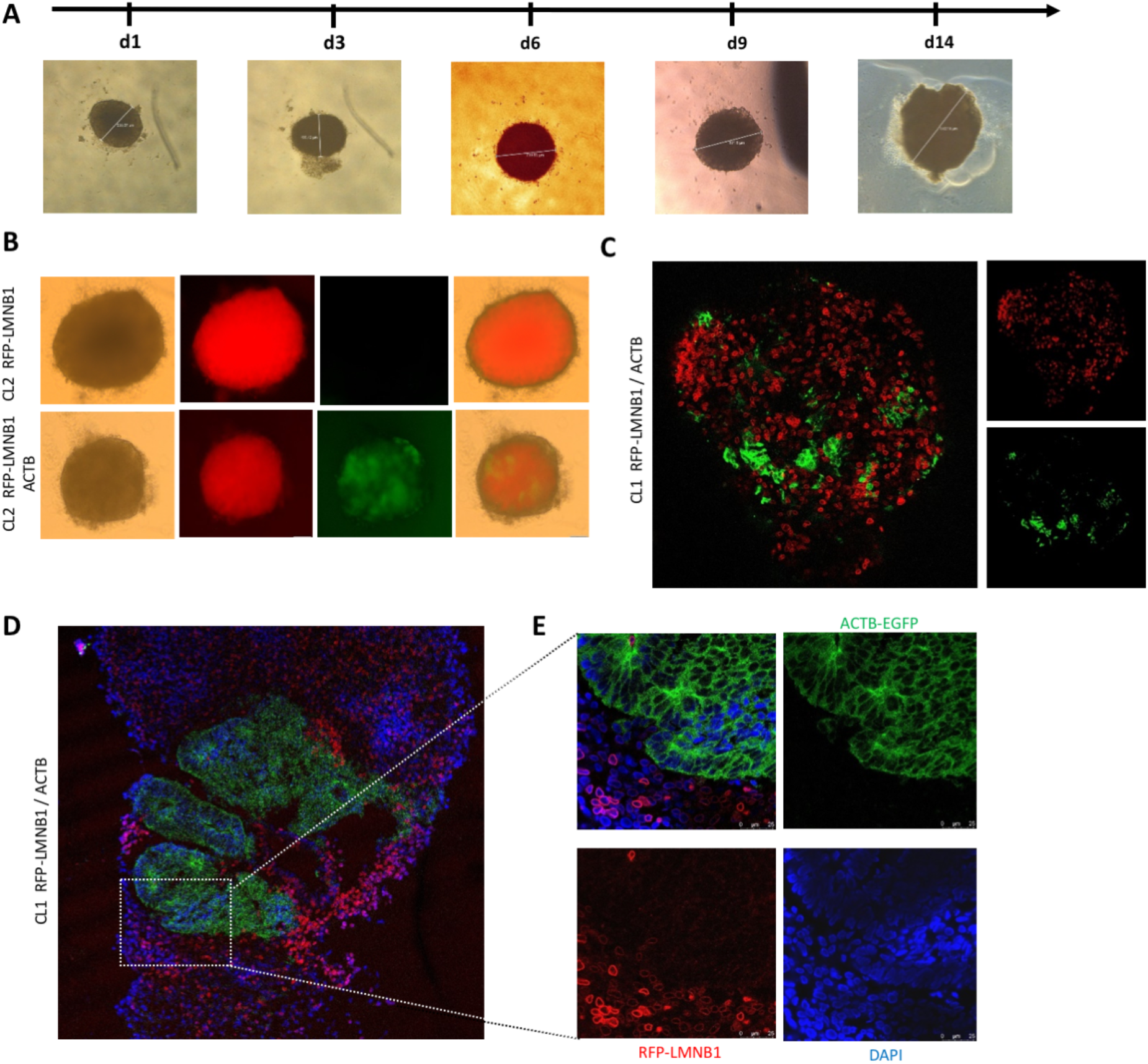
Gene edited RFP-LMNB1 and related mixed organoids to validate the concept of mixed labeled organoids. **(A)** Gene-edited RFP-LMNB1 hiPSCs were used to differentiate EBs, which demonstrates normal development including a roundish shape, translucent edges (>day 8) and increasing diameter. **(B)** The presence of the endogenous label in the RFP-LMNB1 and mixed (50 % RFP-LMNB1 and 50 % ACTB-GFP) EBs is illustrated by fluorescence widefield microscopy (10x objective). **(C)** Confocal microscopy images using 20x objective demonstrated the uneven pattern-like distribution within the CO of the RFP-LMNB1 and the ACTB-GFP cells lines. **(D)** Confocal microscopy images of 50% RFP-LMNB1 and 50 % ACTB-GFP COs stained for DAPI validated the endogenous label and the area-like differentiation within the very same organoid. **(E)** Pattern-like distribution are evident in ACTB-GFP/RFP-LMNB1 cerebral organoid slices (100x objective).

## Discussion

Mixed cerebral organoids (COs) are hybrid tissues derived from two different hiPSC lines. In this work, we demonstrate the feasibility of the mixed organoid approach by developing COs from different mixtures of gene-edited hiPSCs with an endogenously expressed ACTB-GFP, an RFP-LMNB1 label, or without label. These COs exhibited normal growth comparable to non-edited COs and they maintained their fluorescence signal during differentiation and cultivation at high endogenous levels, which are detectable using regular fluorescence microscopy available in most cell culture labs. Mixed COs developed tube-like cortical structures including a cortical plate-like layer and a (sub-) ventricular zone surrounding a lumen indicating successful development of brain-like features and thus suggest that the genetic background is negligible when generating mixed COs. We further demonstrated that individual cells within the mixed COs could be matched to their respective origin according to their specific label. Mixed COs developed patch-like patterns with distinct areas originating from the two hiPSC lines, respectively.

Mixed COs are a promising approach because of the current limitation of normal COs that exhibit high inter-organoid variability **(Quadrato et al., 2017), (Kelava and Lancaster, 2016)** complicating the comparison among themselves, e.g. of a patient-derived CO and a healthy control CO. The specific labeling of hiPSCs with different markers as shown in this work overcomes this limitation as it allows the combination of two (or more) cell lines to differentiate a hybrid organoid containing both, the patient- and control cells within the very same organoid, which allows a more reliable and direct comparison of both conditions. This comparison can include cell to cell analyses, but as a consequence of the patch-like differentiation, the comparison of patient-derived neuronal clusters with control clusters is feasible as well. An ideal experiment would include a patient-derived hiPSC and its isogenic control in order to limit effects caused by the genetic background. A different genetic background might lead to rejection between the different innate immune systems of the cell lines preventing successful merging. However, for the different hiPSC lines used in this study, such effects could not be observed. In addition, rejection could also be prevented by treatment with respective immune system suppressors.

The use of a fluorescent marker enables the detection of changes on a cellular level in live organoids due to morphological alterations or cell death. As COs are developing live tissues, the experiment might also benefit from the fluorescent marker concerning the precise time point for downstream assays, e.g. the determination of the precise age of a CO when a mutation causes structural changes within the organoid in order to schedule drug treatment experiments. To obtain the best results, the selection of a suitable fluorescent marker is crucial: the protein to be fluorescently labeled should be of high abundance, expressed in almost all cell types, and should not correlate with the disease under investigation as large markers might affect the stability or cleavage of a protein **(Bukhari and Müller, 2019)**. To further increase the reliability of the experiment, vice versa labeled hiPSCs and derived organoids are indicated. A clear advantage is the direct visibility in the organoid with no need of further staining (e.g. with myc tag staining). Long-time observations of developing organoids are possible using vibratome-sections, which also allow drug treatment analyses, for example. Furthermore, the fluorescent label enables easy establishment of a labeled hiPSC line, as the fluorescent activated cell sorting allows the fast assessment of the cell line purity.

Another strength of the mixed CO approach is its straightforward integration into downstream analytics. As an example, scRNAseq analyses from COs benefit from the pre-labeled hiPSCs in the respect that expression profiles from the patient cells can be separated from that of the control by including the respective marker as a discriminator in the data analysis pipeline. While many replicates had to be studied in the past in order to achieve sufficient statistical power, the mixed CO approach is capable of significantly reducing the number of biological and technical replicates.

Future studies investigating human disease relevant mutations are indicated to completely unravel the potential of mixed cerebral organoids. In addition to this, the approach is of relevance for personalized medicine. For example, a patient suffering from a psychiatric disorder is currently treated using a trial-and-error technique in order to find the effective medication. In future, these patients might profit from the mixed organoid method. Using low invasive blood sampling peripheral blood mononuclear cells are derived and used for the reprogramming into hiPSCs. Subsequent differentiation of mixed cerebral organoids (using a (primary) relative sample as control) and pharmacological testing of hundreds of agents on mixed CO might significantly shorten the search for the appropriate treatment.

## Acknowledgements

This work was supported by funding from DFG (MU3525/3-2), Mercur (Pr-2016-0010) and BMBF (OrganSARS, 01KI2058). We are thankful to the fluorescence activated cell sorting (FACS) service of the Ruhr University Bochum (Dr. Marcus Peters, Experimental Pneumology, Ruhr-University, Bochum). We want to further give our thanks to Prof. Dr. Andreas Faissner and Dr. Ursula Theocharidis (Department of Cell Morphology and Molecular Neurobiology, Ruhr University, Bochum) for instructing us with and giving us access to their nucleofector.

## Author Contributions

Conceptualization, L.B., L.G.V., N.H., and T.M.; Methodology, L.B., L.G.V., and N.H.; Investigation, L.B., L.G.V., N.H., D.M.; Writing, L.B., L.G.V., N.H., and T.M.; Supervision, T.M.; Funding Acquisition, T.M.

---

**Supplementary Figure 1.**
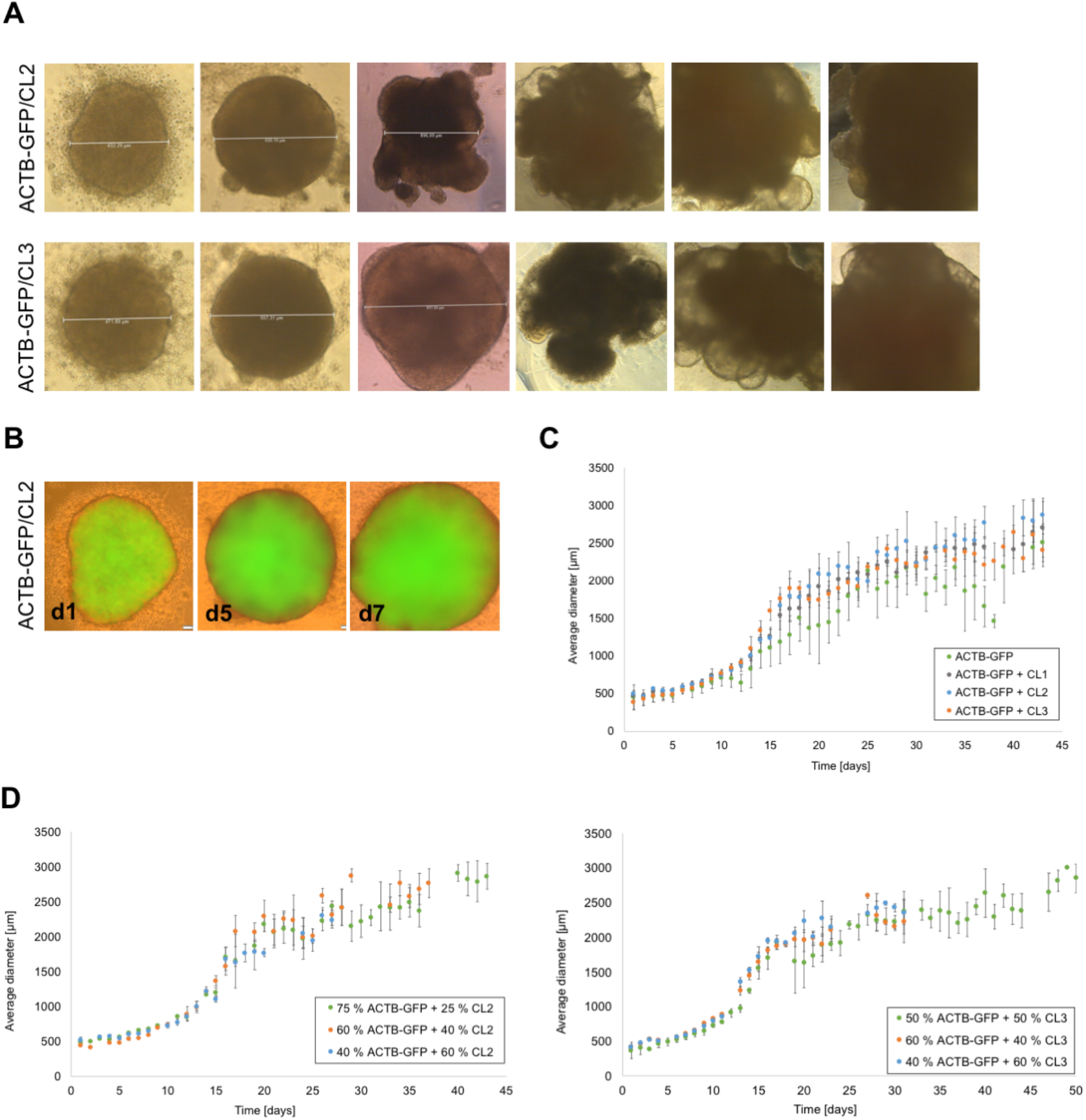
Development of mixed cerebral organoids using CL2 and 3. Cerebral organoids were generated from ACTB-GFP gene-edited hiPSCs mixed with hiPSC lines CL2 or CL3 in different ratios. **(A)** Light microscopy imaging revealed successful formation of EBs with defined edges and a round shape. Translucent edges and the formation of neuroepithelial buds after Matrigel embedding (day 12) was detectable for the mixtures. **(B)** Fluorescence microscopy demonstrated the endogenous label in EBs seeded from 75 % ACTB-GFP + 25 % CL2 hiPSCs. **(C)** A significant difference in size development between COs seeded from different ratios of ACTB-GFP and CL1/CL2/CL3 hiPSCs was not observable. **(D)** Growth of mixed EBs (different mixtures of ACTB-GFP and CL2 or CL3) starting at ~500 μm to 2500 μm after 40 days of cultivation was not different to the growth of pure ACTB-GFP or unlabeled control samples. Error bars correspond to the standard error from at least three biological replicates.

